# Optimization of human papillomavirus-based pseudovirus techniques for efficient gene transfer

**DOI:** 10.1101/2020.04.30.070573

**Authors:** Timra D. Gilson, Ryan T. Gibson, Elliot J. Androphy

## Abstract

Human papillomavirus (HPV) L1 and L2 capsid proteins self-assemble into virions capable of efficiently packaging either its 8 kilobase genome or non-viral DNA. The ability of HPV capsids to package non-viral DNA makes these a useful tool for delivering plasmids to study proteins of interest in a variety of cell types. We describe optimization of current methods and present new protocols for using HPV capsids to deliver non-viral DNA thereby providing an alternative to DNA transfection. Using keratinocyte generated extracellular matrices can enhance infection efficiency in keratinocytes, hepatocytes and neuronal cells. Furthermore, we describe a suspension-based efficient technique for infecting different cell types.

## Introduction

Human papillomaviruses (HPV) are double-stranded non-enveloped DNA viruses that infect keratinocytes of stratified squamous epithelia. The HPV genome is roughly 8,000 base pairs in size, encoding 8 proteins required for completion of the viral life cycle. The two late genes encode L1 (major) and L2 (minor) capsid proteins, which form an icosahedral lattice around the HPV genome. HPV capsids efficiently package non-viral DNA, which presents the possibility of utilizing HPV capsids as a gene delivery vehicle. The practicality of utilizing HPV capsids as gene transfer vehicles could impart advantages over traditional transfection-based gene transfer methods. Lentiviral, adenoviral, and other viral based gene transfer methods are referred to as transduction, however publications utilizing HPV capsids to deliver non-viral DNA into cells use the term infection, which is maintained herein.

To exploit HPV capsids for either HPV genome or gene transfer, a method must be established to easily produce virus. Our intent was to compare the published processes of harvesting HPV capsids and simplifying and optimizing the infection protocol. Early studies using HPV virions required the laborious and inefficient isolation of virions from primary tumors ^1^, xenografts, and rafted keratinocytes, which yield very limited quantities of virus. Over-expression of L1 and L2 proteins was later found to be sufficient for self-assembly of viral particles ^2^. While L1 alone can form native capsids, DNA encapsidation requires L2 proteins ^3,4^. Codon optimization of L1 and L2 DNA sequences was found to increase protein expression by about 100-fold, enabling production of high titer stocks ^5–7^. Currently, the most common L1/L2 expression plasmid is a bi-cistronic 12 kb plasmid (p16SheLL or p16LLw) with an IRES between HPV-16 L1/L2 genes ^8^. The 12 kb plasmid size enables copious production of viral L1/L2 proteins but cannot be encapsidated itself due to its size. The two most common cell lines utilized for viral production are HEK293 TT ^9–15^ and HEK293 TTF cell lines ^16^. Both HEK293 TT and HEK293 TTF cell lines express large and small SV-40 T antigen to drive robust plasmid expression, but HEK293 TTF cells also overexpress the furin protease needed for viral capsid maturation ^17^. When HPV genomes are co-transfected along with p16SheLL into HEK293 TT cells, the genomes are packaged into capsids and these virions are termed quasivirions (QsV); similarly when non-viral DNA up to 8 kb is packaged, virions are called pseudovirions (PsV) ^6,9,10,18^. The development of a robust system for production of HPV QsV and PsV has provided a valuable tool for studying HPV infection and biology, however, our initial studies revealed inefficient infection in both primary and immortal cell lines, which represent the most biologically relevant systems for studying HPV infection.

We examine and compare current methods used in HPV QsV/PsV production and infection, as well as identify improved conditions to mediate delivery of plasmids via PsV into multiple cell types. Additionally, we describe a new infection method for HPV PsV that efficiently mediates gene transfer in difficult to transfect cell lines. Taken together, we find that HPV PsV represent a highly adaptable and efficient tool for studying HPV biology and as an underutilized gene delivery vehicle.

## Results

Several immortal non-cancerous keratinocyte cell lines such as N/TERT, HaCaT and NIKS as well as primary human foreskin keratinocytes (HFK) are commonly utilized for examination of the HPV life cycle. We first sought to determine efficiency and optimize conditions for infection of these cell lines. We utilized PsV packaged with a plasmid encoding mCherry red protein and calculated efficiency of viral infection as percentage of red fluorescent cells, indicating nuclear viral uptake and gene expression. We first titrated the amount of virus by infecting HEK293 TT cells with 1-10 μL mCherry PsV to identify what volume of PsV resulted in 90-95% mCherry positive cells. Two days post infection, 100% infection rates were observed with 5 μL and 10 μL samples and indicated the presence of excess PsV (Fig. 1A), while 3 μL PsV resulted in 90-95% mCherry positive cells. All subsequent infections used 3 μL mCherry containing PsV. Multiple independently produced mCherry PsV batches yielded consistent infectivity, suggesting equivalent titers are produced between batches. PsV infections of N/TERT and HaCaT cells were dramatically reduced compared to HEK293 TT and HeLa cells (Fig. 1B). Low infectivity of N/TERT cells is consistent with previously published results ^19^.

**Figure 1.**
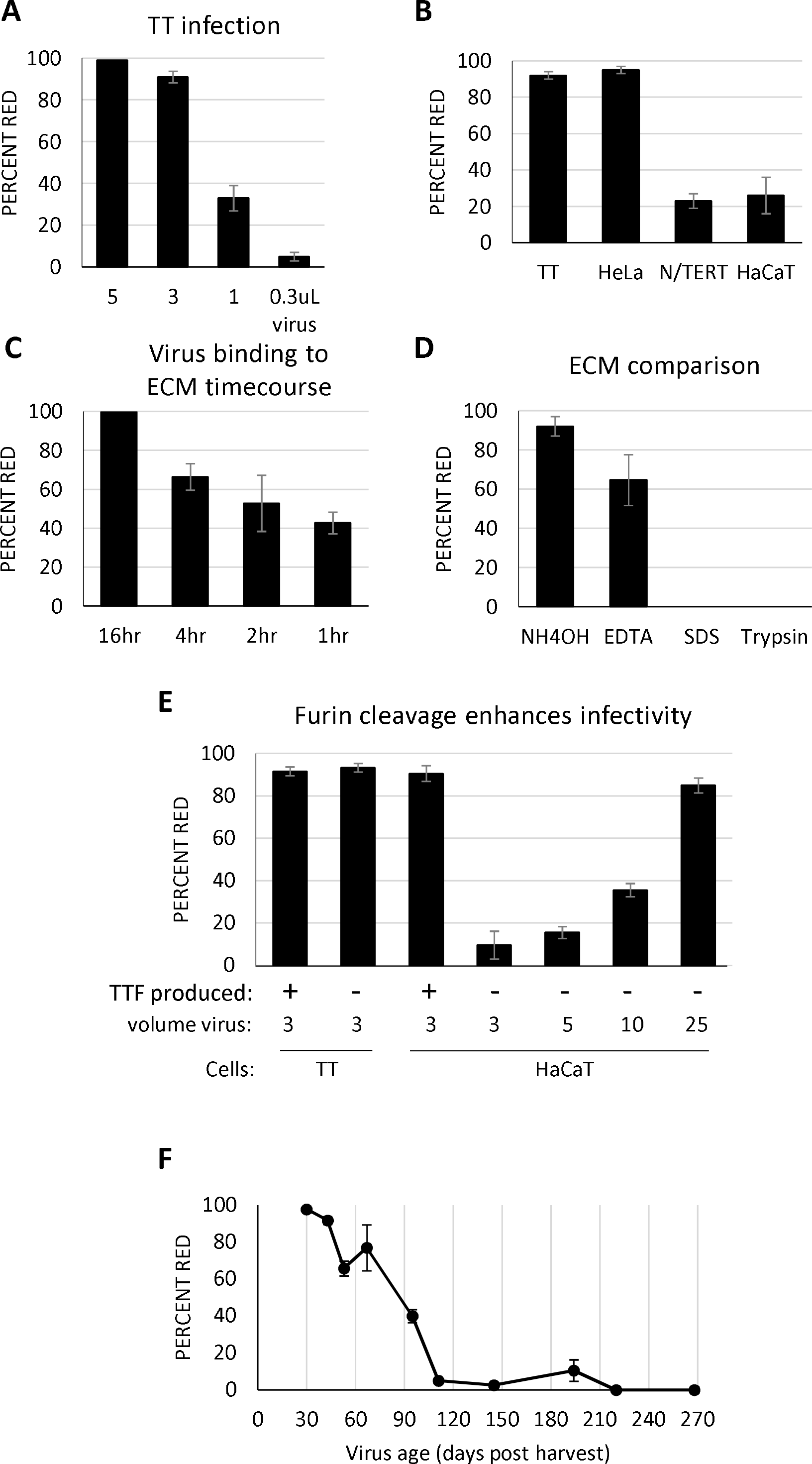
Furin cleavage and ECM promote HaCaT infection. **(A-F)** Data are shown as percent mCherry red fluorescent cells 48 hours post-infection. **(A)** HEK293 TT (TT) cells were plated in a 24-well format. The following day, indicated volumes (μL) of mCherry PsV were added to cells. **(B)** Indicated cell lines were cultured overnight before infection with 3 μL mCherry PsV. **(C)** PsV binding to ECM time-course. HaCaT cells were grown to confluence and NH_4_OH buffer added to remove cells and leave ECM. mCherry PsV was added to ECM for indicated times before unbound virus was removed. **(D)** Comparison of different ECM methods. HaCaT cells were grown to confluence then removed with either NH_4_OH, 10mM EDTA, 0.1% SDS, or 0.05% Trypsin. mCherry PsV was incubated with ECM overnight and HaCaT cells were added next day. **(E)** HEK293 TT (TT) produced PsV are not as infectious as HEK293 TTF (TTF) produced PsV. TT cells were incubated with 3 μL TT made mCherry PsV overnight. HaCaT cells were grown to confluence, stripped with NH_4_OH/Triton-X buffer and incubated with TT produced mCherry PsV overnight. The following day, HaCaT cells were added. **(F)** mCherry PsV is stable for one month. mCherry PsV stored for the indicated number of days at 4°C were added to HaCaT generated ECM overnight. Media was changed the next day and HaCaT cells added. All experiments were performed at least 3 independent times and values shown as mean ± SEM.

To attempt to increase infection rates of these immortal keratinocyte cell lines, we sought to identify factors influencing infection. During infection HPV binds to and penetrates the milieu of proteins secreted by cells known as the extracellular matrix (ECM) before interacting with a cell surface receptor. ECM binding is thought to induce conformation changes in L1/L2 that promote virus internalization. HPV has been found to most efficiently bind ECM derived from epithelial cells ^11,20^, and some labs have employed the use of epithelial cell derived ECM for studies of HPV infection. To our knowledge, a direct comparison of infection efficiency using PsV pre-bound to ECM versus PsV directly applied to cells adhered to tissue culture plastic has not been published. Therefore, we evaluated whether pre-binding PsV to HaCaT ECM would increase infection rates of epithelial cells. HaCaT cells were grown to confluence to generate ECM, then cells removed using NH_4_OH/Triton-X 100 ^20^ and mCherry PsV allowed to adhere to ECM for the indicated times. After removal of media containing unbound virus, HaCaT cells were added and examined for red fluorescence two days later. One-hour incubation of PsV with ECM increased HaCaT infection efficiency more than 3-fold over PsV not pre-bound with ECM (Fig. 1C). Overnight incubation of PsV with ECM resulted in nearly 100% infection.

Another published method to generate ECM is to remove cells with EDTA. As these two different methods likely strip different subsets of proteins, we sought to compare the efficiency of these methods in promoting infection. Both NH_4_OH and EDTA methods generated ECM able to bind PsV and promote infection of HaCaT cells (Fig. 1D). Two additional methods for generating ECM failed to promote infection: 20 seconds in 1 % SDS, and 8 minutes in 0.05 % Trypsin (Fig. 1D), consistent with previous experiments using HPV-11 virions ^11^. Given that we find the NH_4_OH method promotes the highest infection rates, removes cells in roughly 15 - 60 seconds versus 10 min - 2 hrs using EDTA (current paper, ^11,15^) and EDTA generated ECM requires repeated washing to remove all cells and/or mitomycin C treatment to prevent growth of any remaining cells ^15^, all subsequent ECMs described herein followed the NH_4_OH method.

HEK293 TTF cells were utilized for PsV production. Derived from HEK293 TT cells, HEK293 TTF cells stably overexpress the furin protease needed for viral capsid maturation ^16,17^. Furin cleaved PsV have been reported to have increased infectivity compared to non-cleaved virus ^16,20–22^. Several other cell lines have been tested to produce PsV including green monkey CV-1 cells ^2^ and insect SF9 cells ^23^, but literature most frequently cites the use of HEK293 TT cells ^9–15^. Therefore, we compared infection rates of PsV generated in HEK293 TT to HEK293 TTF. While HEK293 TT cells are equally infectable with both HEK293 TT and HEK293 TTF produced PsV, HaCaT cells required more HEK293 TT produced PsV to generate infection rates equivalent to HEK293 TTF produced virus (Fig. 1E). This is in agreement with previous findings that HEK293 TT cells showed equal infection rates ^16^ and psga-745 cells showed higher infection rates with furin cleaved PsV ^20^. Given the increased infectivity, HEK293 TTF cells were used for all subsequent PsV production. PsV stored at 4^°^C is stable for one month, after which infection rates start to decrease (Fig. 1F).

Next, we sought to examine whether pre-binding PsV to ECM would similarly improve N/TERT infection. We tested HaCaT as well as N/TERT generated ECM, however both failed to promote N/TERT infection (Fig. 2A). Given the different growth requirements of HaCaT (DMEM/5% FBS) and N/TERT cells (serum-free KSFM), it was possible that factors present in KSFM are inhibitory to infection and/or FBS factors are stimulatory to infection. Direct comparison of individual factors in the different media could not be performed as KSFMs formulation is proprietary. To determine the effects of media on infection, ECM attached PsV were incubated in different media prior to infection, then media changed to KSFM before adding N/TERT cells to ECM/PsV. FBS containing media such as F-media, E-media and DMEM-FBS all stimulated N/TERT infection, but media lacking FBS (e.g. Optimem, DMEM) did not (Fig. 2B). FBS contains many factors including growth factors, hormones, and cytokines, which are removed through incubation with charcoal. ECM incubated with this stripped FBS did not promote infection nearly as well as normal FBS (Supplementary Fig S1). Similarly, horse serum did not efficiently stimulate infection (Supplementary Fig S1). As N/TERT cells prefer KSFM media, we sought to determine if adding 5% FBS to KSFM would increase infection, and were surprised that no stimulation was observed (Fig. 2C). However if FBS was instead added to basal KSFM lacking the standard additives Bovine Pituitary Extract (BPE) and hEGF, increased infection was detected (Fig. 2C). As both E- and F-media contain hEGF while promoting high infection rates, we suspected BPE is inhibitory to PsV infection. We found that adding BPE to F-media significantly decreased infection rates (Fig. 2C). Therefore, PsV infection of N/TERT cells was enhanced by using ECM and FBS-containing media, while BPE inhibited PsV infection. N/TERT cells showed negligible infection with HEK293 TT produced PsV compared to HEK293 TTF generated virus (Fig. 2D).

**Figure 2.**
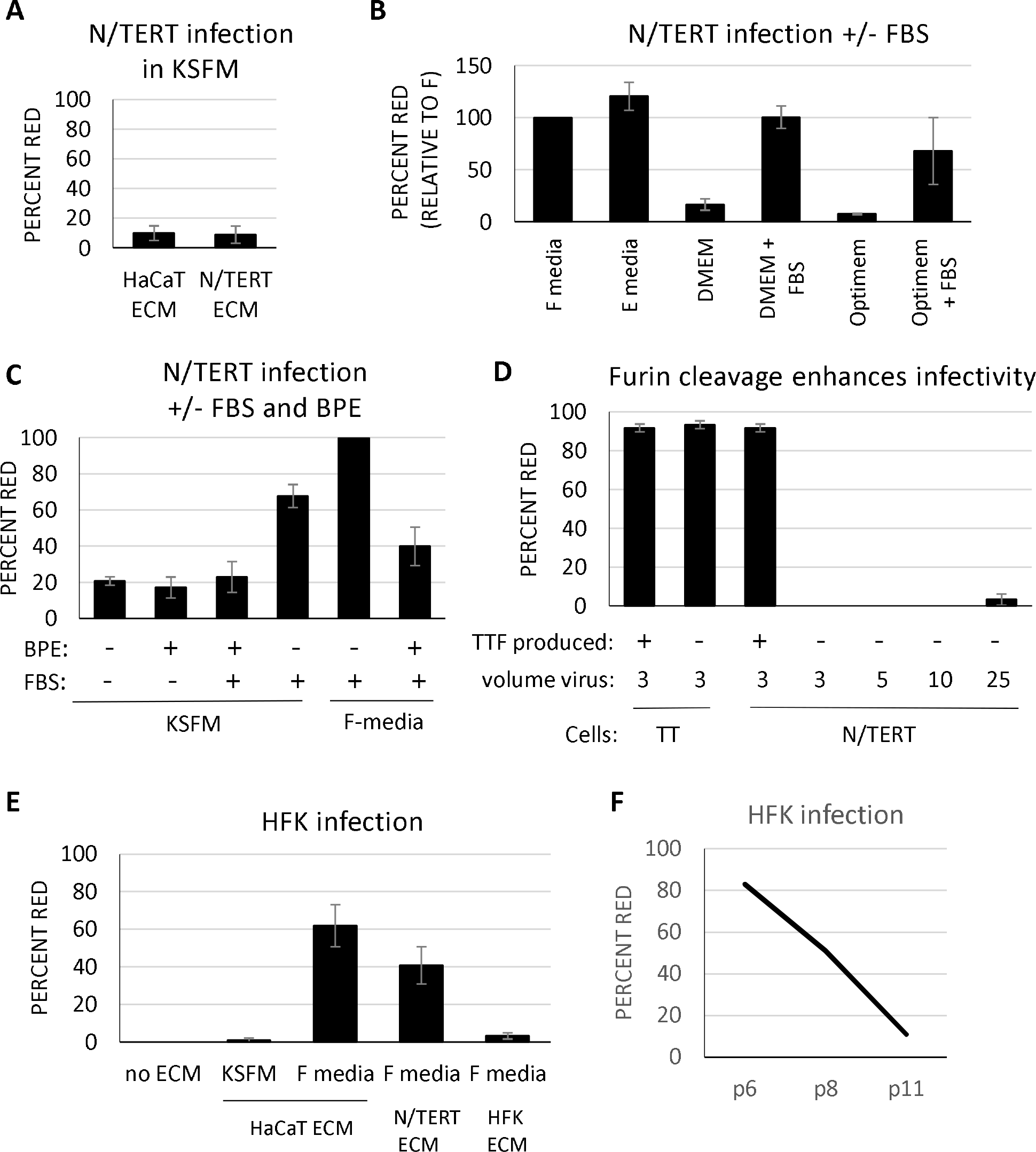
N/TERT and HFK infection optimization. **(A)** ECM does not promote infection in KSFM. HaCaT or N/TERT cells were grown to confluence, stripped with NH_4_OH buffer, and incubated with mCherry PsV overnight in KSFM before adding N/TERT cells in fresh media. (**B)** FBS promotes infection. N/TERT infections were performed as in A, except PsV was incubated in the indicated media. (**C)** BPE inhibits infection. N/TERT infections were performed as in A except the indicated media were used. (**D**) Furin cleavage promotes efficient infection. HEK 293TT (TT) cells were infected with 3 μL TT or TTF produced PsV, and N/TERT cells were incubated with 3 μL TTF PsV or 3, 5, 10, or 25 μL TT produced PsV. For (**B-F)**, following overnight incubation of PsV with ECM, media were removed and replaced with KSFM and N/TERT cells added. (**E)** Serum and ECM promote infection. ECMs generated from the indicated cells were incubated with mCherry PsV in the indicated media overnight, then media was changed to KSFM and HFKs added. (**F)** High passage HFKs show reduced infectivity. HaCaTs ECM was incubated with mCherry PsV in F-media overnight, then HFKs of the indicated passage numbers were added. For all parts, two days following infection, infection efficiency was determined as percent of red cells. All experiments were performed at least 3 independent times and values shown as mean ±SEM.

Primary HFKs are the best model for HPV replication but are not efficiently infected by PsV ^14,19,24^. Given that N/TERT cells were derived from HFK cells, we tested our optimized protocol for N/TERT infection with HFK. Indeed, no infection was seen without ECM, while PsV pre-bound with HaCaT ECM in F-media resulted in roughly 50% HFK infection, and only 1% observed using KSFM media (Fig. 2E). Primary cell lines such as HFKs only replicate for a limited number of passages, and proliferation slows as passages increase ^25^. The number of passages undergone by HFKs at the time of infection influences infection efficiency. Early passage HFKs (<6 passages) are highly infectable, but extended culturing results in decreased infection rates (Fig. 2F).

In the presence of FBS, HaCaT generated ECM significantly increases infectivity of HaCaT, HFK, N/TERT keratinocytes. To determine if these modified conditions can be utilized as a universal infection protocol, additional cell types typically used in PV studies were examined. Cervical cancer human C33A cells and Bovine (B)PV replication competent mouse mammary epithelial C127 cells are utilized in PV infection assays. Both displayed low infection rates in the absence of ECM. PsV pre-bound to HaCaT ECM in the presence of standard growth media DMEM + 5% FBS dramatically increased infection rates (Fig. 3A). Similarly, the infection rate of replicating mouse fibroblast J2 3T3 cells is nearly 100 % in presence of ECM. J2 3T3 cells are co-cultured with NIKS and CIN612-9E cells to prevent differentiation, however, these are non-replicating due to mitomycin-c treatment. Given that cells must undergo mitosis for packaged DNA to enter the nucleus ^26^, we anticipated mitomycin-c treated J2 cells (J2 MC) would be resistant to infection, and indeed this was observed (Fig. 3B). Therefore, infection of NIKS and CIN612-9E cells can be performed in the presence of co-cultured mitomycin-c treated J2 3T3 cells.

**Figure 3.**
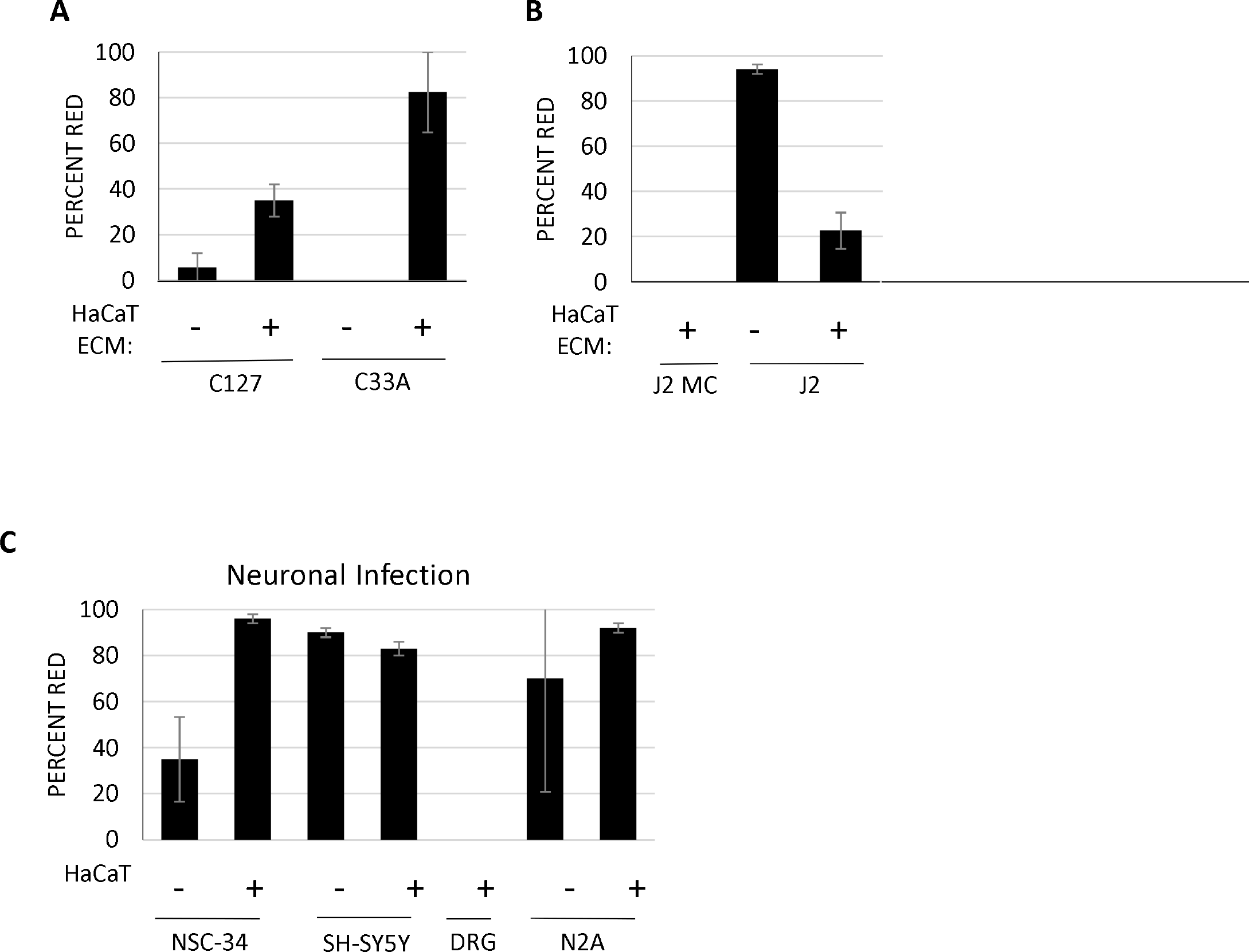
HaCaT ECM promotes infection of different cell types. **(A-B)** Cells plated prior to PsV exposure were not efficiently infected (no ECM lanes). HaCaT generated ECM was incubated with mCherry PsV overnight in DMEM/FBS before media was replaced in (**A)** with C33A and C127 cells, and (**B)** with J2 cells treated with mitomycin-C overnight or left untreated before adding cells to PsV bound ECM. (**C)** HaCaT ECM was incubated with mCherry PsV overnight before NSC-34 mouse motor neuron like, SH-SY5Y human neuroblastoma, and DRG mouse primary dorsal root ganglia neurons (non-replicating) were added. Infection efficiency was calculated as percent of red fluorescent cells two days post-infection. All experiments were performed at least 3 independent times and values shown as mean ± SEM.

Exposure to HPV occurs *in vivo* through epidermal or mucosal contact, and there is no evidence for hematogenous spread. Nonetheless, PsV infect a wide array of replicating cells including neuronal cells ^14^. We speculated that this could be improved using our ECM protocol. Immortal neuronal cell lines such as the neuroblastoma N2A and SH-SY5Y as well as motor neuron hybrid NSC-34 cells were found to be highly infectable (Fig. 3C). This highly efficient infection rate of SH-SY5Y (>80%) with PsV is remarkable considering transfection of these cells is notoriously difficult with rates between 10-30% transfection efficiency (personal observation, and Lipofectamine 3000 product page(https://www.thermofisher.com/us/en/home/brands/product-brand/lipofectamine/lipofectamine-3000.html#validated)). As expected, primary murine dorsal root ganglion (DRG) cells did not express the mCherry transgene as these do not replicate and mitosis is required for nuclear access of the transgene (Fig. 3C).

*In vivo*, HPV infects keratinocytes on the basement membrane. The ECM of the basement membrane keratinocytes is comprised of a complex assortment of fibrous proteins, glycosaminoglycans, and proteoglycans produced by both keratinocytes and fibroblasts. Therefore, we tested the ability of fibroblast generated ECM to promote infection. J2 fibroblast ECM did not stimulate infection of any tested cell lines (Fig. 4). Similarly, ECM from other epithelial cell lines HEK293 TT and HeLa did not support infection of keratinocytes. ECM derived from keratinocyte cell lines NIKS, N/TERT, and HaCaT cells promoted high infectivity compared to no ECM (Fig. 4). HaCaT ECM overall stimulated the highest infection rate of all ECMs tested.

**Figure 4.**
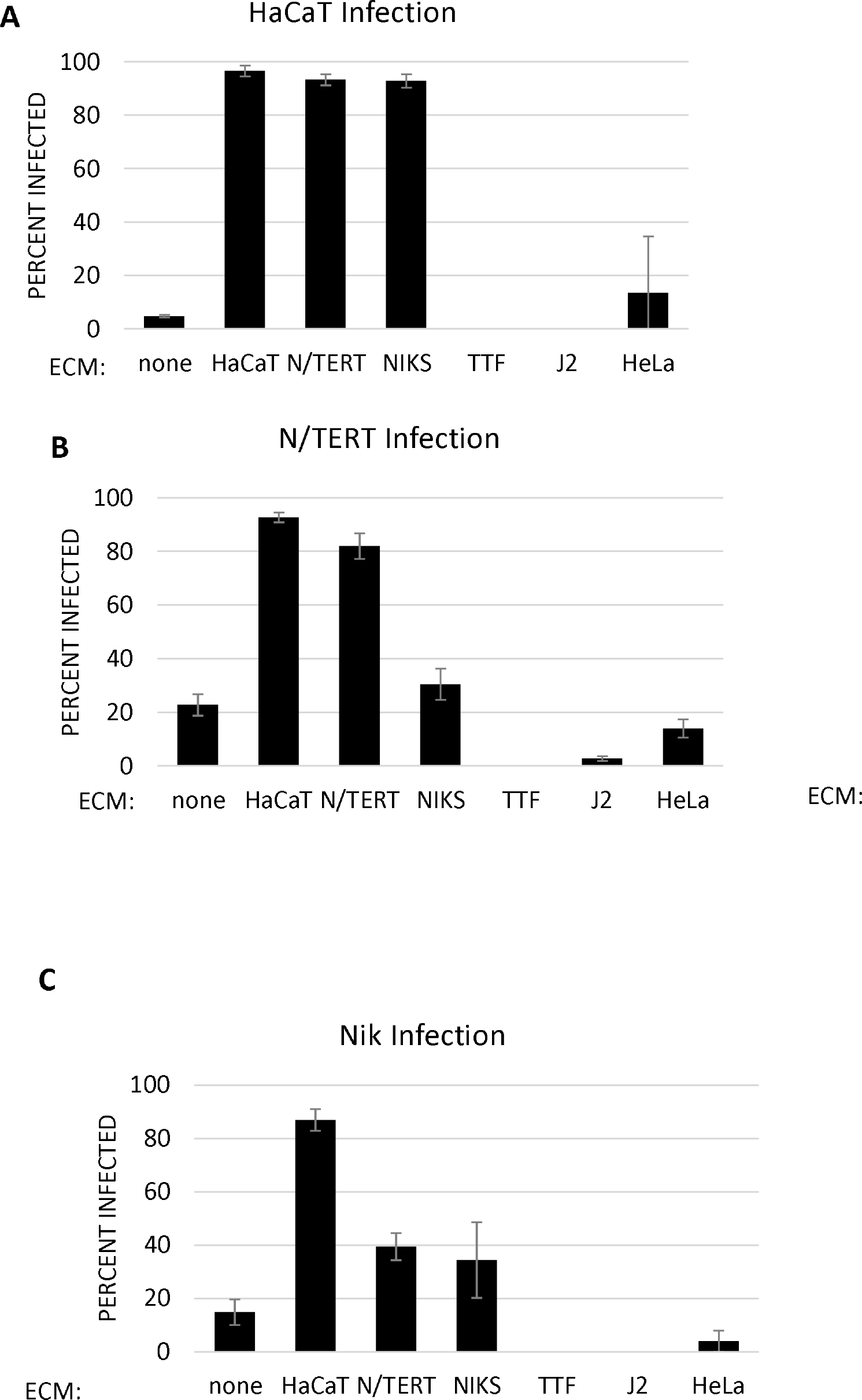
HaCaT ECM promotes the highest infection rate in keratinocyte cell lines. ECMs from the indicated cells at the bottom of each graph were incubated with mCherry PsV overnight in low calcium DMEM/FBS **(A)** or F-media **(B** and **C)**. Unbound PsV were removed, fresh media and HaCaT **(A)**, N/TERT **(B)**, or NIKS **(C)** cells then added. Percent red was calculated two days post-infection. All experiments were performed at least 3 independent times and values shown as mean ± SEM.

Since natural HPV infects basal layer keratinocytes, the HaCaT based ECM was generated using low calcium media (0.03 mM) as this promotes a basal phenotype ^27,28^. HaCaT cells grown in standard DMEM media exposed to high calcium (2 mM) become differentiated, as noted by production of involucrin and phenotypic changes resembling differentiated NIKS cells ^27,28^ (Fig. 5A). While differentiated HaCaT cells have been successfully used for ECM, we wanted to directly compare high and low calcium HaCaT ECM. N/TERT infection showed no difference between the ECMs, however both HaCaT and NIKS cells had higher infection on undifferentiated HaCaT ECM (Fig. 5B).

**Figure 5.**
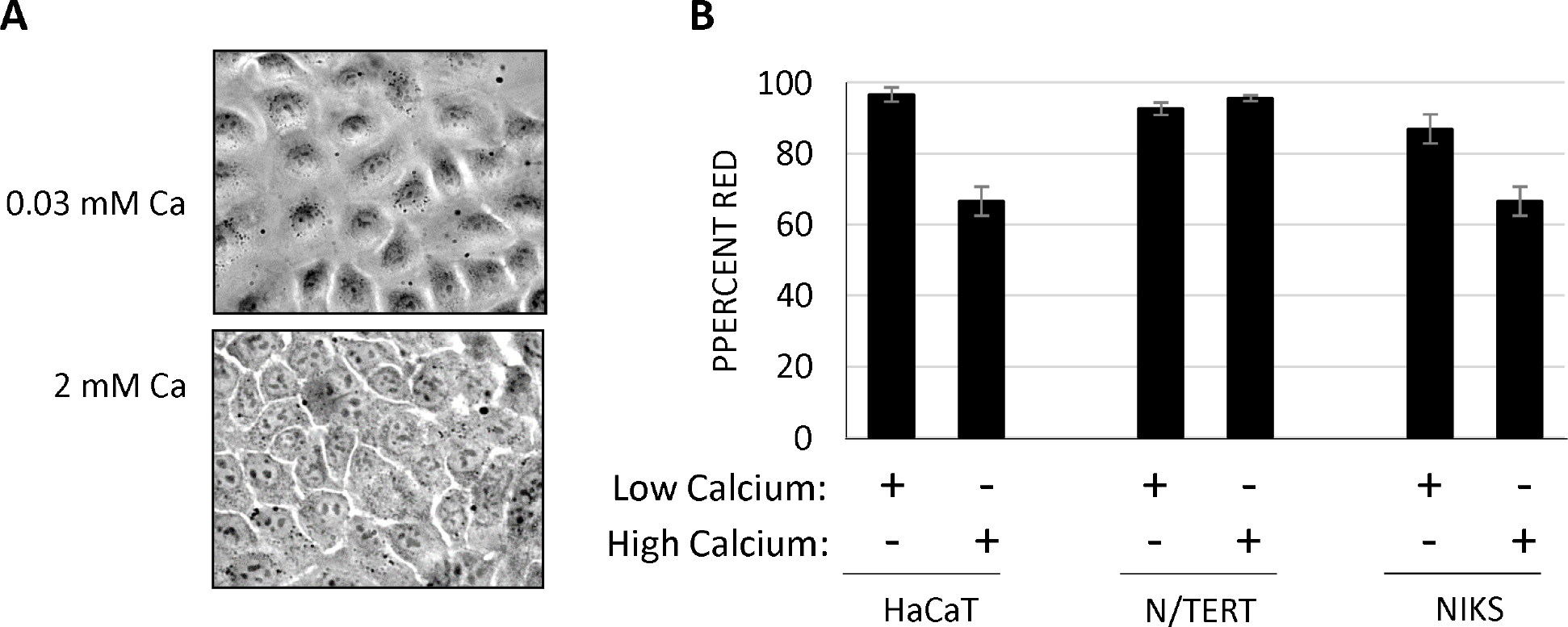
Calcium levels influence quality of HaCaT ECM. **(A)** HaCaT cells grown in low calcium have an extended non-differentiated phenotype. High calcium present in standard DMEM (2mM Ca^2+^) promotes HaCaT differentiation. Pictures are white light images of HaCaT cells in high and low Ca^2+^ media at 40x. (**B)** ECMs generated from HaCaT cells grown in high or low calcium media were incubated with mCherry PsV overnight before indicated cells were added. Percent red was calculated two days post-infection. All experiments were performed at least 3 independent times and values shown as mean ± SEM.

The established model of HPV infection involves interactions of the viral capsid proteins with cellular factors to mediate internalization and retrograde trafficking of the virus within the host cell (Valdano et al., 2019). This process occurs in adherent cells with established polarity and vectoral trafficking. We speculated that increased cell surface exposure may enhance the efficiency of HPV PsV infection. We piloted trials of suspension-mediated infection (SMI) in which trypsinized cells are exposed to virus immediately before plating (Fig. 6A). SMI infection efficiency was higher or equivalent to cells plated prior to infection (Fig. 6B). N/TERT and Huh-7 cells with low efficiency of infection without ECM were efficiently infected with PsV using SMI. HepG2 cells were uninfectable even in the presence of various ECM but nearly 100% susceptible to infection by SMI. Representative images of SMI infected cells are shown in Fig. 6C with differential interference contrast white light images on left and mCherry infected cells on right. Intriguingly, SMI efficiency in various cell lines could be greatly enhanced based on the use of different media. For example, N/TERT cells infected in KSFM with mCherry PsV showed low infection efficiency using SMI, however when used on N/TERT cells in F-media, efficiency increased from roughly 10% to nearly 80% (Fig. 6D). Media also influenced SMI efficiency in HepG2 and Huh-7 cells, with E-media producing the highest infection rates in both cell lines (Fig. 6E-F). N/TERT cells are able to proliferate in serum containing media and while this induces morphological change, these cells reverted within 30 minutes of placement into serum-free KSFM (Fig. 6G).

**Figure 6.**
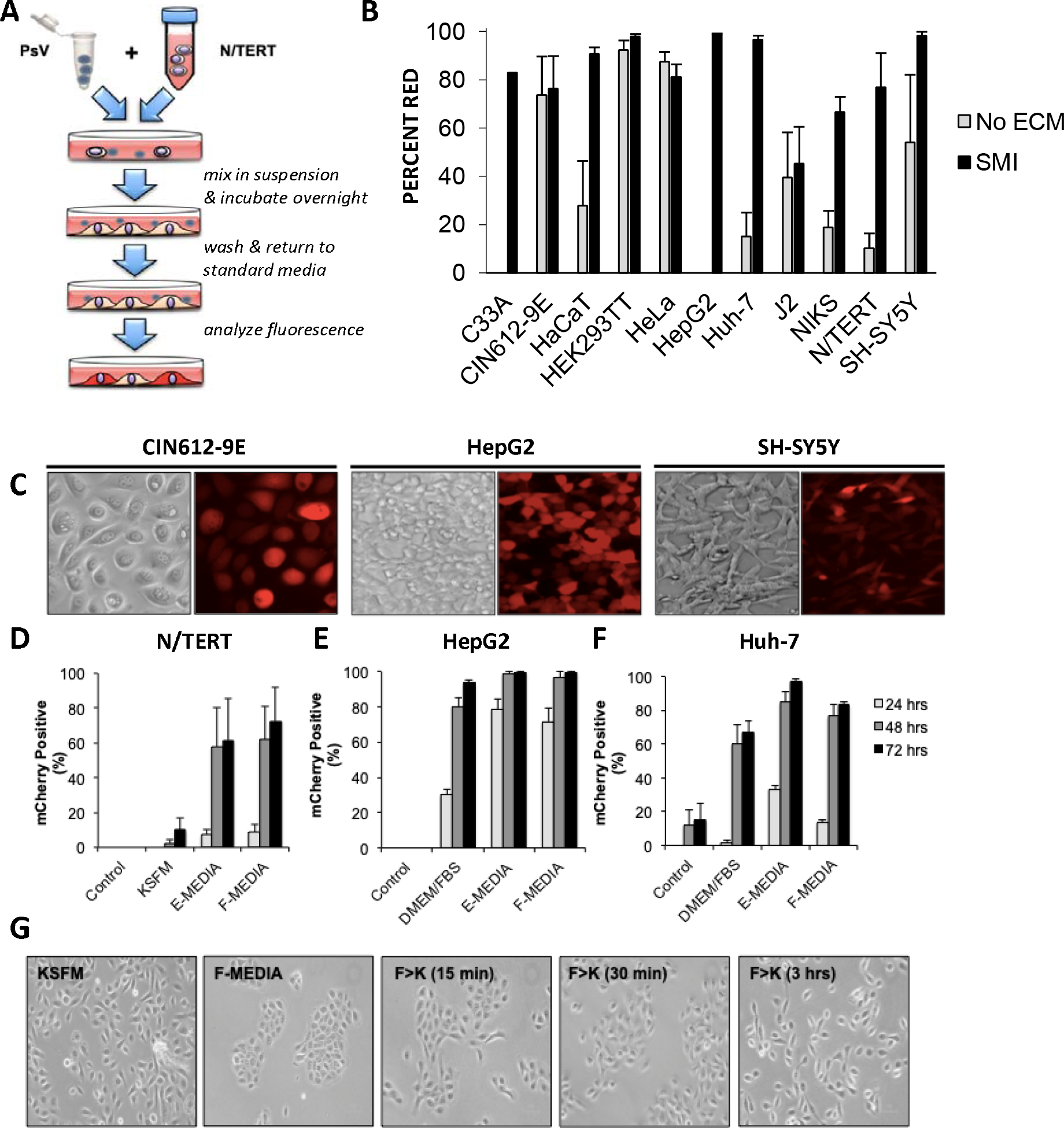
SMI mediated infection. (**A**) Schematic representation of the SMI technique. (**B**) Summary comparison of cell infection efficiency at 72 hrs post-infection with mCherry PsV using media found most efficient for infecting each cell type. Cells were plated without ECM (PP) prior to infection (grey), or infected by SMI (black). Data are percent of cells positive for mCherry fluorescence. (**C)** Immunofluorescence of CIN612-9E, HepG2 and SH-SY5Y cells infected with mCherry PsV using SMI. DIC image on left and fluorescent mCherry images on right. (**D**) N/TERT, **(E)** HepG2, or **(F)** Huh-7 cells were infected with mCherry PsV without ECM (control), or using SMI in different media and fluorescence analyzed at 24, 48, and 72 hrs. All experiments were performed at least 3 independent times and values shown as mean ± SEM. (**G)** Morphology of N/TERT cells observed after plating cells into either KSFM (panel 1) or F-media (panel 2), then N/TERT cells 15 min after changing F-media to KSFM (panel 3), 30 min (panel 4), and 3 hrs (panel 5).

Given that we have demonstrated efficient infection of notoriously difficult to transfect cell lines and that PsV capsids can encapsidate plasmid DNA up to 8 kb in size, we sought to determine if PsV infection via SMI can be used as a transfection alternative with plasmids other than pmCherry. Using SMI, HEK293 TT, N/TERT and SH-SY5Y cells were infected with either mCherry, HPV-31 V5-E2, or HA-εCOP PsV. Two days post-infection, cells were lysed and immunoprecipitated with an antibody to V5 or HA. All three cell types were successfully infected and expressed either HPV-31 V5-E2 or HA-εCOP (Fig. 7A-C). Therefore, SMI constitutes an employable method to mediate efficient plasmid delivery and an alternative to transfection methods. While SMI and ECM based methods both efficiently infect N/TERT cells, we compared these methods for heterologous protein expression. N/TERT cells were infected with V5-E2 PsV either by SMI or ECM method, then two days post-infection cell lysates were immunoblotted with V5 antibody. Remarkably, SMI produced greater V5-E2 protein expression (Fig. 7D).

**Figure 7.**
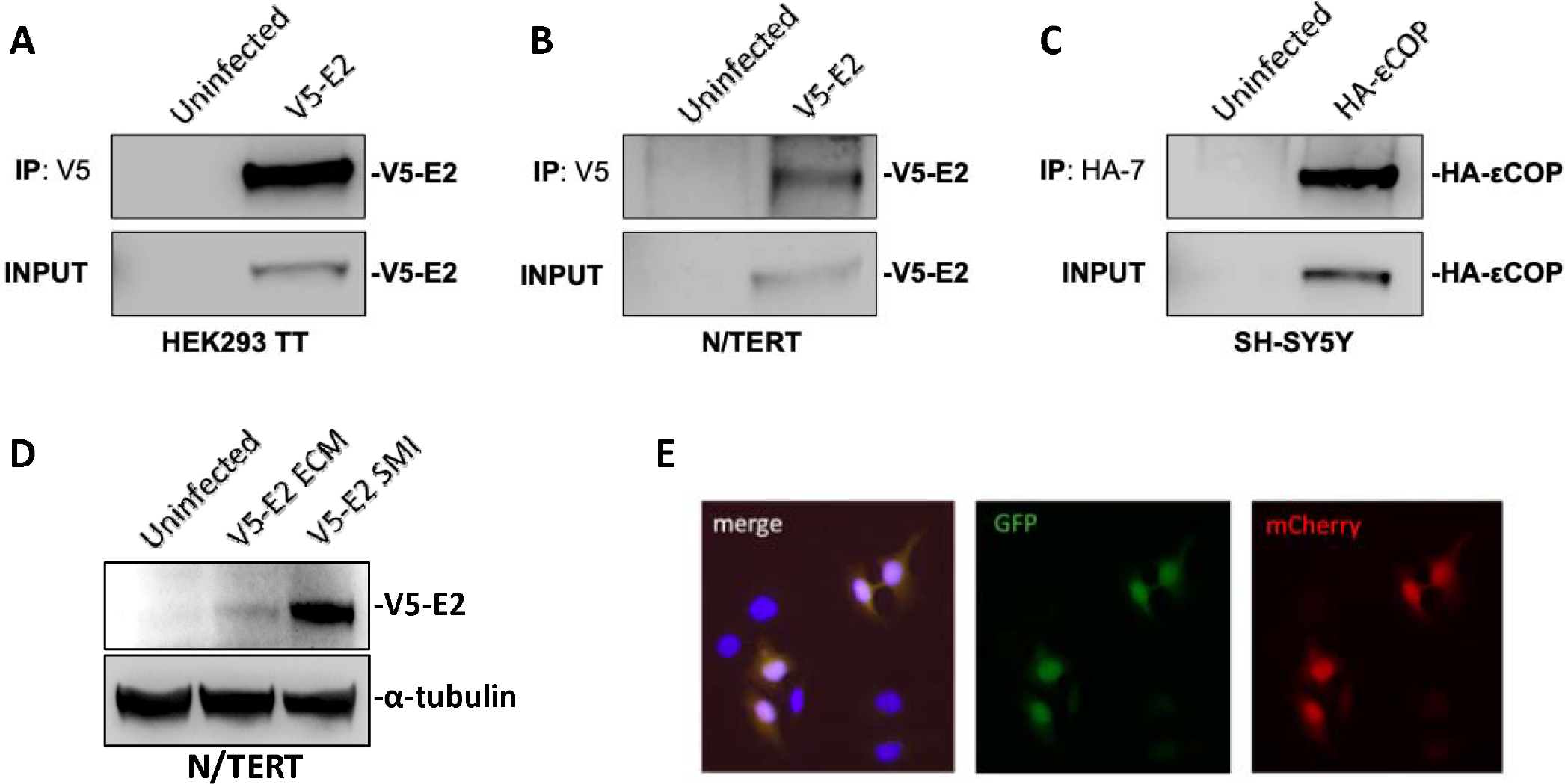
PsV can package multiple DNAs for delivery. **(A)** HEK293 TT or **(B)** N/TERT cells were infected with HPV-31 V5-E2 PsV for two days, lysed and immunoprecipitated with rabbit anti-V5 antibody, then immunoblotted with mouse anti-V5 antibody. (**C)** SH-SY5Y cells were infected with HA-εCOP PsV for two days, lysed, and immunoprecipitated with mouse 12CA5A1 anti-HA antibody and immunoblotted with mouse HA-7 anti-HA. (**D)** N/TERT cells were infected with V5-E2 PsV either by SMI or ECM methods. Two days following infection, cells were harvested and immunoblotted with V5 and Tubulin antibodies. (**E)** HaCaT cells were co-infected with GFP and mCherry PsV.

Oftentimes when cells are transfected, more than one plasmid is utilized and therefore it is advantageous if cells take up both plasmids. To examine the efficiency by which differently packaged PsV infect the same cell, we performed co-infections of HaCaT cells with mCherry and GFP packaged PsV. As shown in Fig. 7E, cells were either dual infected with mCherry and GFP PsV, or remained uninfected. Therefore, introduction of multiple plasmids into cells using PsV co-infections are equivalent to co-transfection.

A summary of all cell lines examined for PsV infection is given in Table 1.

**Table 1:**
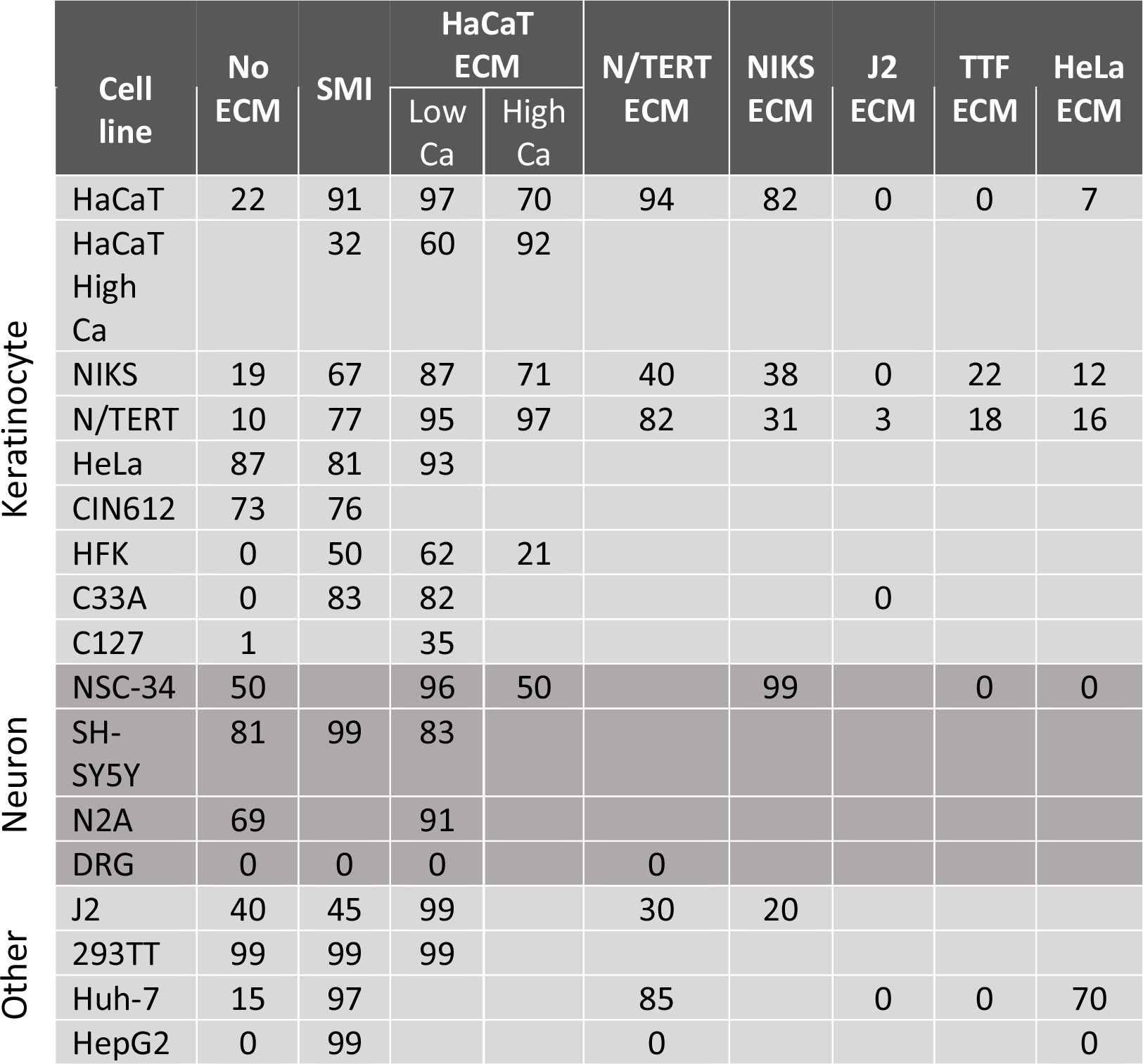
Summary of infectivity rates of various cell lines (left) utilizing no ECM, SMI, or different ECMs (top)

## Discussion

Production of HPV capsids using overexpression of codon optimized L1 and L2 proteins provides an inexpensive, less tedious method of isolating virus. While HEK293 TT and HeLa cells have high infection rates, the keratinocytes of interest for examining HPV infection all have lower infectivity. Using HPV pseudovirions (PsV) with a packaged mCherry reporter gene, we tested existing methods and describe two protocols that result in at least 50% infection of replicating cells.

One protocol combines four steps: 1) using NH_4_OH generated keratinocyte extracellular matrix (ECM) _20_, 2) pre-binding of PsV to ECM ^11^, 3) adding serum ^12^, and 4) using furin cleaved virus ^16^. While HEK293 TT, HeLa, and CIN612-9E cells infect well without added ECM, HaCaT, N/TERT, NIKS, and primary HFK cells require all four conditions for optimal infection efficiency. This may reflect that these non-transformed cells recapitulate natural infection, in which HPV first attaches to the ECM and basal keratinocytes and the L1 and L2 proteins undergo conformational changes and proteolytic cleavages before entering the cell (reviewed in _29_).

Furin proteases cleave after Rx(R/K)R motifs, two of which are present in the N-terminus of L2 capsid protein. Post-translational processing is required for packaged DNA to escape endosomes and proceed to the Trans Golgi Network ^17,30^. Production of PsV/QsV in furin overexpressing HEK293 TTF cells is advantageous over standard HEK293 TT produced virus, as HEK293 TTF cells allow for greater capsid maturation. The ECM is a complex assortment of proteins, proteoglycans, glycoproteins, and collagen ^31^. Different cell types secrete unique ECM factors, and our study agrees with others ^11^ that keratinocyte (HaCaT, NIKS, N/TERT) ECM provides the highest infection rates. Interestingly, while HeLa cells are readily infected without ECM, these cells are a poor source of ECM for infection of other keratinocyte cell lines. Nonetheless, HeLa derived ECM renders Huh-7 hepatocytes highly susceptible to infection, whereas Huh-7 ECM does not. The buffer used for cell removal when generating ECM likely strips off different subsets of polysaccharides and proteins. With the NH_4_OH/Triton method, collagen, heparin sulfate, sulfated glycosaminoglycans (sGAG), laminin, and fibronectin are preserved upon extraction ^32,33^. We have successfully generated ECM using EDTA stripping methods established in other labs, but we prefer the NH_4_OH method due to the shorter incubation time as well as the ability to circumvent repeated pipetting to remove all cell soma. Serum contains factors required for infection as N/TERT and HFK cells propagated in keratinocyte serum free media (KSFM) demonstrated negligible PsV uptake. With serum such as found in DMEM/FBS or F-media, nearly 100% infection was observed. Growth factors present in FBS likely contribute to stimulating virus binding as charcoal stripped FBS did not stimulate infection.

ECM and SMI based infection methods both achieve high infection rates, but each method provides distinct advantages. The simpler, faster SMI method involves simply combining PsV and trypsinized cells together in suspension at the time of plating. The resulting protein expression can be higher than ECM method (Fig. 7D). However, the limitation of SMI is the amount of virus utilized before toxicity occurs. For the longer more involved ECM method, much more virus can be utilized before cell toxicity is noted, and is the preferred method for cells growing in serum free media. The requirement of serum during infections can transiently change the morphology of cells normally propagated in serum free media. While we have noted this altered morphology reverts quickly back to normal in SMI infected N/TERT cells upon changing back to serum free media (Fig. 6G), this may not be true for all cells. Pre-binding PsV to ECM in the absence of cells has the advantage of avoiding serum exposure to serum-sensitive cells.

It is unclear why there is a difference in protein expression between SMI and ECM methods. One theory is that the amount of PsV entry is relative to surface area, such that the increased cell surface of floating cells in SMI method provide more viral attachment points compared to the ECM method where only the bottom of cells are virus exposed. If this is the case, we would expect large HaCaT cells to have higher infection rate than small HEK293TT cells, but instead we observed the opposite in the absence of ECM (Fig 1). Similarly we would expect SMI infection rates to be around twice as high as ECM infection rates, but that is not the case for HepG2 cells which are completely resistant to infection with ECM but become ∼100% infectable using the SMI technique. These observations imply that surface area does not correlate with PsV infection.

HPV infection occurs following microabrasions exposing the basement membrane and polarized adherent basal keratinocytes to HPV ^34,35^. Following interactions of the viral capsid proteins with ECM, apical cellular proteins, and receptors, the HPV capsid is internalized and undergoes retrograde endosomal trafficking toward the nucleus, where it resides within vesicles until breakdown of the nuclear envelope ^36^. The requirement for ECM interactions and cellular polarity are challenged in the context of the SMI method. SMI mixes cells and PsV in suspension and results in efficient infection of multiple cell types, including those cells resistant to transfection or found uninfectable with or without ECM. This is potentially explained by exposure of cells to PsV in suspension enabling greater cellular surface access or exposure to factors on the cell undersurface not normally accessible until mitosis as cells release from the culture plate. The observation that non-dividing DRGs cannot be infected by PsV supports the concept that cell division is required for nuclear uptake.

Transfection as a method for delivering plasmids into cells for genome or protein overexpression has other limitations. The presence of transfected DNA in the cytoplasm is detected by cGAS, resulting in activation of the antiviral cGAS-STING pathway ^37,38^. This may reduce gene expression at both transcriptional and translational levels. Variable transfection efficiency between different cell types makes consistent delivery and expression of genes challenging, as many cell lines demonstrate low transfection efficiency. Both cost and toxicity of some reagents highlight additional drawbacks with the use of transfection. These issues associated with transfection based methods are not limited to protein overexpression but also manifest when studying HPV biology. Upon infection, L1/L2 encapsidated viral genomes undergo retrograde trafficking from endosomes through the Trans Golgi Network (TGN) prior to entering the nucleus ^30^. By avoiding cytoplasmic exposure, HPV does not expose the viral genome to innate antiviral cytosolic DNA sensors. Accurately studying HPV biology necessitates the avoidance of triggering pathways that would otherwise not be stimulated.

Additionally, the number of HPV genomes introduced to each cell by transfection is much higher than what occurs during natural infection. Taken together, the limitations associated with transfection based approaches in the study of HPV biology and for gene delivery highlight the importance of developing more efficient methods of DNA delivery.

Multiple viruses currently in use as gene transfer vehicles (Adenovirus, adeno-associated virus, lentivirus, and baculovirus) require the gene-of-interest to be subcloned into a special viral packaging vector before infectious virus can be produced. The ability of HPV capsids to package standard mammalian expression vectors could impart an advantage over traditional viral gene transfer methods. We have demonstrated that PsV can package pCI and pcDNA3 based vectors, although other vectors may or may not be suitable for PsV packaging. One drawback of HPV PsV based gene delivery is the limitation to dividing cells, whereas adenovirus, adeno-associated viruses and lentiviruses are able to infect non-replicating cells. In conclusion, HPV PsV-mediated infections provide a method to study HPV infection and a versatile technique for efficient delivery of plasmids up to 8 kb in size into a variety of cell types.

## Methods

### Cell lines

C33A (D. Lowy), C127 (D. Lowy), HeLa (D. Lowy), J2 3T3 (J2) (H. Green), HepG2 and Huh-7 (H. Guo), HEK293 TT (J. Schiller), HEK293 TTF (R. Roden), N2A (A. Burghes), NSC-34 (N. Cashman), and SH-SY5Y (ATCC) were cultured in DMEM + 10% FBS. CIN612-9E (L. Laimins) cells were maintained in E-media with mitomycin-C treated J2 cells, primary human foreskin keratinocytes from newborn discards (HFK) and N/TERT (I. Morgan) in KSFM, HaCaT (N. Fusenig) cells in low calcium DMEM/5% FBS, and spontaneously immortalized keratinocyte (NIKS, B. Allen-Hoffman) were passaged in F-media with mitomycin-C treated J2 cells. Trypsinization of cells for passaging was performed by incubation with 0.05% Trypsin-EDTA solution for 2-10 minutes.

### Plasmids

p16sheLL expressing codon optimized HPV-16 L1 and L2 capsid proteins separated by an IRES for PsV production ^8^ and pmCherry and pfwb (GFP) ^9^ for quantitation of viral infection were provided by J. Schiller. pCIneo-V5-HPV31-E2 was cloned with a V5 oligo (Nhe-V5-Afl-F CTAGC atgGGCAAGCCTATCCCAAACCCTCTGCTCGGACTGGACAGTACT C and Nhe-V5-Afl-R TTAAG AGTACTGTCCAGTCCGAGCAGAGGGTTTGGGATAGGCTTGCCcat G) and inserting into Nhe/Afl II digested pCIneo. Codon optimized HPV31-E2 (from A. McBride) was amplified from pcDNA3-FLAG-31E2 with primers (Mlu-31E2-F GATC acgcgt ATGGAGACCCTGagcc and Not-31E2-R GTAC GCGGCCGC TTAAATGGTC) and inserted into Mul/Not sites within pCIneo-V5. pcDNA3-HA-εCop was cloned through PCR amplification of εCOP cDNA isolated from HaCaT cells with primers Nco-hεcop-F GATC CCATGG CGCCTCCG and hεcop-Xho-R GTAC CTCGAG TCAGGCGCTGGGA, digestion with Nco/Xho, and insertion into pcDNA3-HA.

### PsV production

PsV stocks were made using a modified protocol combining the original PsV method (https://ccrod.cancer.gov/confluence/display/LCOTF/Home) with an optimized furin cleavage method ^16^. HEK293 TTF cells were plated at 1 ⨯ 10^7^ cells in 15 cm plates for next day co-transfection with 20 μg p16sheLL and 20 μg pmCherry using Lipofectamine 2000 and incubated overnight. Media was changed and non-essential amino acids (Gibco) added the next day. Two days post-transfection, cells were collected, washed 3x with PBS, and gently resuspended in 1.5 cell pellet volume equivalents viral maturation buffer (0.5% Triton X, 10 mM MgCl_2_, 5 mM CaCl_2_, 25 mM (NH_4_)_2_SO_4_, PBS), then incubated at 37^°^C for 2 days in siliconized microcentrifuge tubes. Lysates were spun at 7000xg for 5 minutes and soluble viral stock stored at 4°C that retained high infectivity for up to one month (Fig. 1A).

### Extracellular matrix (ECM) production and Infection

HaCaT, J2, HEK293 TTF, HeLa, NIKS, HFK, or N/TERT cells were grown to confluence. Cells were washed in PBS and treated with ECM buffer containing either NH_4_OH or EDTA to remove cell bodies but leave residual ECM on the plate. For NH_4_OH based ECM (Day 2012), PBS was removed from rinsed cells and 0.3 mL ECM buffer (170 mM NH_4_OH, 0.5% Triton, PBS) added per well for one minute. ECM buffer was removed, ECM washed 3x with PBS, complete removal of cells was visually confirmed via light microscopy, and 0.5 mL media plus 3 μL mCherry PsV added. After overnight incubation, unbound PsV was removed, media changed and cells plated. Two days post infection, cells were visualized for red fluorescence to determine infection efficiency. For the EDTA isolated ECM, PBS was removed and 0.5 mL EDTA buffer (10 mM EDTA in PBS) was incubated with cells for 10 minutes at 37°C. A few cells are removed with gentle tapping while the majority remain in the periphery and were removed by vigorously pipetting.

### Suspension-mediated infection (SMI)

SMI was performed by mixing 2 ⨯ 10^4^ cells and 3 μL PsV in suspension at the time of plating, allowing PsV to bind to cells in suspension prior to adhesion to plates and in the absence of ECM, then the cells were incubated overnight at 37^°l^C. The following day, media containing unbound virus was removed and intracellular red fluorescence visualized at 24, 48 and 72 hrs.

### Immunoprecipitation (IP) and Immunoblotting

HEK293 TT, N/TERT and SH-SY5Y cells were infected with either mCherry, HPV-31 V5-E2, or HA-εCOP PsV. Two days after infection, cells were lysed in 0.5% NP-40, 150 mM NaCl, 20 mM Tris (pH 7.5) with protease inhibitor cocktail and rotated for one hour at 4^°^C with benzonase. Following centrifugation, soluble lysate was collected and IP performed by incubation of lysates with Protein A/G slurry and either rabbit anti-V5 (Cell Signal Technologies) or mouse 12CA5A1 anti-HA antibodies. Beads were washed in lysis buffer, boiled in 2X Protein Sample buffer, run on SDS-PAGE gels, and transferred onto 0.45 μM PVDF membranes (Millipore) by semi-dry transfer. Membranes were blocked in 5% non-fat milk/PBS/0.1% Tween-20 then incubated overnight at 4^°^C with designated primary antibodies. ECL (Amersham) chemiluminescence substrates were used for protein detection using an ImageQuant LAS 4000 system (GE Healthcare).

### Statistical analysis

All experiments were repeated a minimum of three times and data are expressed as mean ± standard error of the mean (SEM).

## Acknowledgements

We appreciate the generosity of Alison McBride (NIAID), John Schiller and Chris Buck (NCI) for providing plasmids and the cited sources of the cell lines we used. John Schiller, Patricia Day and Nathan Fons kindly offered helpful comments on our manuscript. This research was supported by the National Cancer Institute R01CA058376 to EJA. National Institute of Allergy and Infectious Diseases T32AI007637 and T32AR062495 to TG. The content is solely the responsibility of the authors and does not represent the official views of the NIH.

## Author Contributions

T.D.G. and R.T.G. performed experiments. T.D.G., R.T.G. and E.J.A. conceptualized the study, designed experiments and interpreted data. T.D.G., R.T.G. and E.J.A. wrote and reviewed the manuscript.

---

**S Figure 1.**
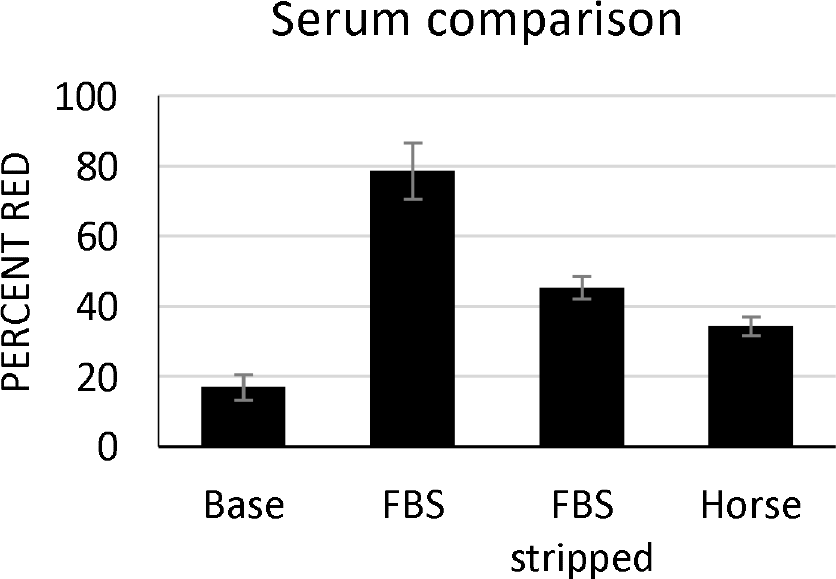
Serum type effects infection rate. N/TERT cells were grown to confluence, stripped with NH_4_OH buffer, and remaining ECM was incubated with mCherry PsV overnight in media lacking any serum (base) or supplemented with FBS, charcoal stripped FBS, or Horse Serum. N/TERT cells were added next day in fresh F-media and percent red cells was calculated 48 hours post infection. All experiments were performed at least 3 independent times and values shown as mean ± SEM.

**Supplementary Figure S2.**
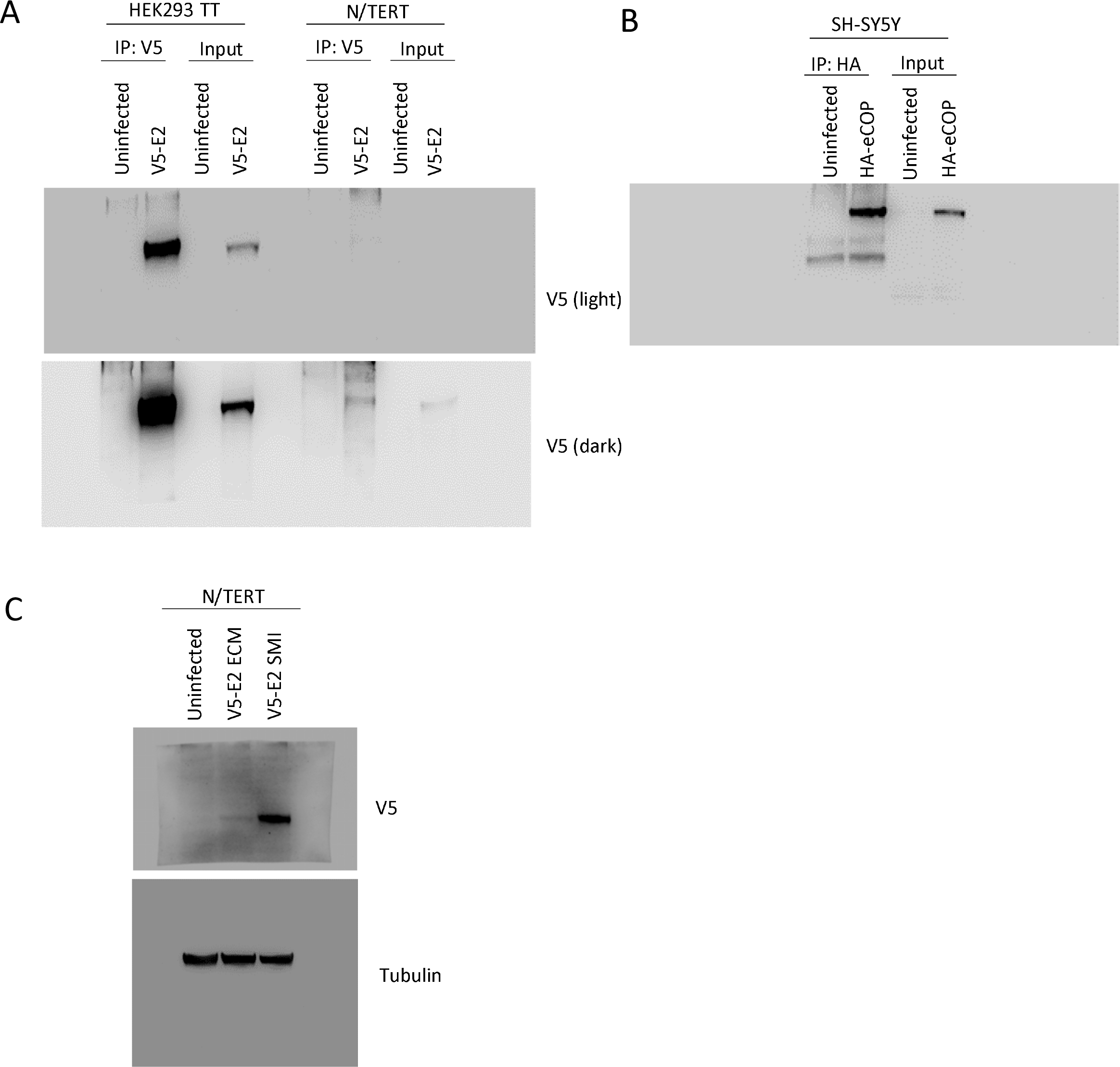
Full length blots for Figure 7. **(A)** Full length blot for Fig. 7A on left (lighter exposure), and Fig. 7B on right (darker exposure). HEK293 TT or N/TERT cells were infected with HPV-31 V5-E2 PsV for two days, lysed and immunoprecipitated with rabbit anti-V5 antibody, then immunoblotted with mouse anti-V5 antibody. **(B)** Full length blot for Fig 7C. SH-SY5Y cells were infected with HA-εCOP PsV for two days, lysed, and immunoprecipitated with mouse 12CA5A1 anti-HA antibody and immunoblotted with mouse HA-7 anti-HA. **(C)** Full length blot for Fig 7D. N/TERT cells were infected with V5-E2 PsV either by SMI or ECM methods. Two days following infection, cells were harvested and immunoblotted with V5 and Tubulin antibodies.

